# Conspicuous candidate alleles point to *cis*-regulatory divergence underlying rapidly evolving craniofacial phenotypes

**DOI:** 10.1101/2020.03.23.003947

**Authors:** Joseph A. McGirr, Christopher H. Martin

## Abstract

Developing a mechanistic understanding of genetic variation contributing to variation in complex craniofacial traits is a major goal of both basic and translational research. Investigating closely related species that evolved divergent feeding morphology is a powerful approach to identify genetic variation underlying natural and clinical variation in human craniofacial phenotypes. We combined whole-genome resequencing of 258 individuals with 50 transcriptomes to identify candidate *cis*-acting genetic variation influencing rapidly evolving craniofacial phenotypes within an adaptive radiation of *Cyprinodon* pupfishes. This radiation consists of a dietary generalist species and two derived trophic niche specialists – a molluscivore and a scale-eating species. Despite extensive morphological divergence, these species only diverged 10 kya and produce fertile hybrids in the laboratory. Out of 9.3 million genome-wide SNPs and 80,012 structural variants, we found very few alleles fixed between species – only 157 SNPs and 87 deletions. Comparing gene expression across 38 purebred F1 offspring sampled at three early developmental stages, we identified 17 fixed variants within 10 kb of 12 genes that were highly differentially expressed between species. By measuring allele-specific expression in F1 hybrids from multiple crosses, we found strong evidence for two *cis*-regulatory alleles affecting expression divergence of two genes with putative effects on skeletal development (*dync2li1* and *pycr3*). These results suggest that SNPs and structural variants contribute to the evolution of novel traits and highlight the utility of the San Salvador pupfish system as an evolutionary model for craniofacial development.

## Introduction

Craniofacial anomalies account for approximately one-third of all birth defects (Gorlin et al. 1990). These include jaw deformities, oral clefts, defects in the ossification of facial or cranial bones, and facial asymmetries. Much of what is known about the developmental genetic basis of craniofacial morphology and function comes from mutagenesis screens and loss of function experiments in model organisms (Hall 2009). These types of studies have been critical to identifying genes essential for craniofacial development and alleles underlying monogenic disease conditions that exhibit Mendelian inheritance. However, screens are biased to detect alleles within protein-coding regions that severely disrupt gene function and are likely to cause lethality at early developmental stages (Nguyen and Tian 2008; Hall 2009). Furthermore, it is now understood that much of the natural and clinical variation in complex traits like craniofacial morphology results from interactions among hundreds to thousands of loci across the genome (Boyle et al. 2017; Sella et al. 2019). Genome-wide association studies (GWAS) have shown that the vast majority of genetic variants affecting complex traits and diseases are within non-coding regions, highlighting the importance of gene regulation influencing trait variation (Hindorff et al. 2009; Maurano et al. 2012; Schaub et al. 2012). Thus, complementary approaches to mutagenesis screens in model organisms are necessary to identify genes that influence craniofacial phenotypes at later stages in development though changes in gene regulation rather than gene function.

One such approach is to harness naturally occurring genetic variation between ‘evolutionary mutants’ – closely related species exhibiting divergent phenotypes that mimic human disease phenotypes (Albertson et al. 2008). Several fish systems have been particularly useful as models for craniofacial developmental disorders because closely related species are often distinguished by differences in morphological traits important for trophic niche specialization, such as the shape and dynamics of jaws and pharyngeal elements (Albertson et al. 2008; Schartl 2014; Powder and Albertson 2016). The process of identifying candidate genes and validating their effect on phenotypic divergence in evolutionary mutants typically involves population genomic analyses, gene expression analyses, GWAS, and functional validation experiments (Bono et al. 2015; Kratochwil and Meyer 2015). Using a combination of these approaches, research in fish systems has shown that the evolution of adaptive craniofacial traits often involve orthologs of genes implicated in human disorders (Albertson et al. 2005; Helms et al. 2005; Roberts et al. 2011; Ahi et al. 2014; Cleves et al. 2014; Lencer et al. 2017; Erickson et al. 2018; Gross and Powers 2018; Martin et al. 2019). Therefore, candidate genes identified in evolutionary mutant models that have orthologs with uncharacterized functions in humans warrant further study into their relationship with development and disease.

Advances in next generation sequencing technologies alongside substantial reductions in the cost of sequencing have made it possible to sequence the genomes of hundreds of individuals and identify millions of single nucleotide polymorphisms (SNPs) and structural variants (SVs) segregating between closely related species. Measuring relative and absolute genetic differentiation (estimated as *Fst* and *Dxy*) between species can reveal diverged regions of the genome that may influence trait development, but these statistics alone are insufficient to identify genetic mechanisms underlying evolutionary mutant phenotypes (Nachman and Payseur 2012; Cruickshank and Hahn 2014). RNA sequencing across multiple developmental stages and tissue types can provide further evidence that differentiated regions influence phenotypic divergence if genes near genetic variants are differentially expressed between species (Whiteley et al. 2010; Poelstra et al. 2014; McGirr and Martin 2018; Verta and Jones 2019). However, this assumes that linked genetic variation within *cis*-acting regulatory elements affects proximal gene expression levels, and does not rule out the possibility of unlinked *trans*-acting regulatory variation binding regulatory regions to influence expression levels (Wittkopp and Kalay 2011; Signor and Nuzhdin 2018). Determining whether a candidate gene is differentially expressed due to *cis*- or *trans*-regulatory divergence is important to identify putatively causal alleles that can be further validated by genome editing or transgenesis experiments.

It is possible to use RNAseq to identify mechanisms of gene expression divergence between parental species by bringing *cis* elements from both parents together in the same *trans* environment in F1 hybrids and quantifying allele specific expression (ASE) of parental alleles at heterozygous sites (Cowles et al. 2002; Wittkopp et al. 2004; Signor and Nuzhdin 2018). ASE occurs when a heterozygous allele within a coding region that is alternatively homozygous in two parental species shows biased expression in F1 hybrids. *Cis-*regulatory divergence is expected when a gene is differentially expressed between species and shows ASE in F1 hybrids; whereas *trans*-regulatory divergence is expected when a gene is differentially expressed and does not show ASE (Wittkopp et al. 2004; Davidson and Balakrishnan 2016; Signor and Nuzhdin 2018). Thus, genes showing signs of *cis*-regulatory divergence that are near differentiated regions of the genome make promising candidates for causal variation underlying evolutionary mutant phenotypes, especially when the same genes show high genetic differentiation between species and are implicated by GWAS. Together, these strategies can target candidate variation with nucleotide level resolution and provide a framework to prioritize variants for functional validation experiments.

Here, we combine whole-genome resequencing, RNAseq, and F1 hybrid allele specific expression analyses to identify candidate *cis*-acting genetic variation influencing rapidly evolving craniofacial phenotypes within an adaptive radiation of *Cyprinodon* pupfishes on San Salvador Island, Bahamas (Fig. 1). This sympatric radiation consists of a dietary generalist species (*C. variegatus*) and two endemic specialist species adapted to novel trophic niches – a molluscivore (*C. brontotheroides*) and a scale-eater (*C. desquamator*; (Martin and Wainwright 2013a)). Nearly all forty-nine pupfish species in the genus *Cyprinodon* distributed across North America and the Caribbean are dietary generalists with similar craniofacial morphology that is used for consuming algae and small invertebrates (Fig. 1A (Martin and Wainwright 2011; Martin and Wainwright 2013b)). The molluscivore evolved short, thick oral jaws stabilized by a nearly immobile maxilla allowing it to specialize on hard-shelled prey including ostracods and gastropods (Fig. 1B). This morphology results in a larger in-lever to out-lever ratio compared with generalists, increasing mechanical advantage for strong biting (Hernandez et al. 2018). The molluscivore is also characterized by a prominent maxillary anteriodorsal protrusion that may be used as a wedge for extracting snails from their shells (Martin et al. 2017). The scale-eater is a predator that evolved to bite scales and protein-rich mucus removed from other pupfish species during rapid feeding strikes (Fig. 1C (St. John et al. 2020)). Scale-eaters have greatly enlarged oral jaws, larger adductor mandibulae muscles, darker breeding coloration, and a more elongated body compared with the generalist and molluscivore species (Martin and Wainwright 2013a).

**Fig. 1.**
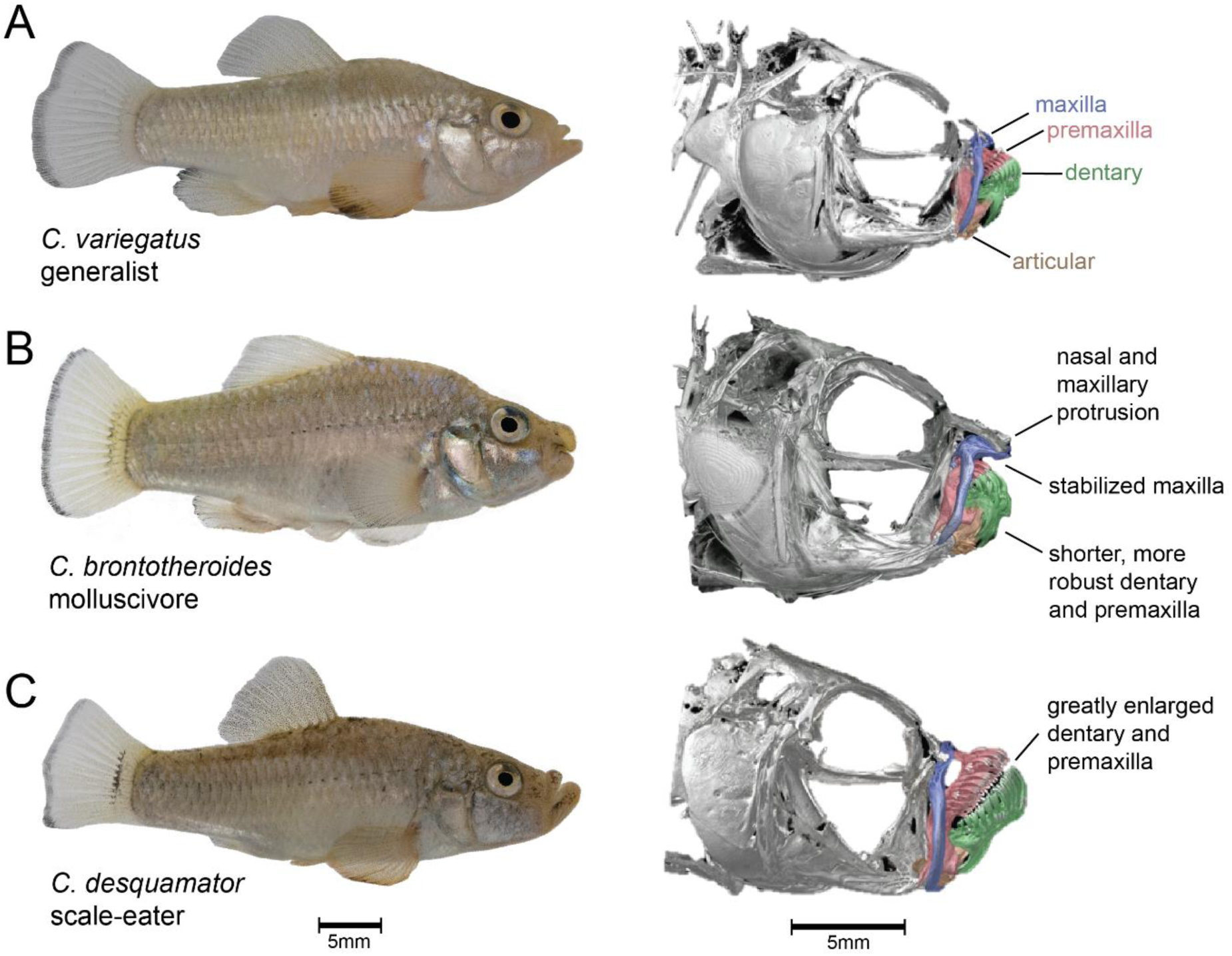
San Salvador Island pupfishes exhibit exceptional craniofacial divergence despite recent divergence times. A) *Cyprinodon variegatus* (generalist), B) *C. brontotheroides* (molluscivore), C) *C. desquamator* (scale-eater). µCT scans modified from (Hernandez et al. 2018) show major craniofacial skeletal structures diverged among species including the maxilla (blue), premaxilla (red), dentary (green), and articular (brown).

Exceptional craniofacial divergence despite extremely recent divergence times and low genetic differentiation between molluscivores and scale-eaters make this system a compelling evolutionary model for human craniofacial developmental disorders. These trophic specialist species rapidly diverged from an ancestral generalist phenotype within the last 10-15k years (Turner et al. 2008; Martin and Feinstein 2014). Molluscivores and scale-eaters readily hybridize in the laboratory to produce fertile F1 offspring with approximately intermediate craniofacial phenotypes between the parents and no obvious sex ratio distortion (Martin and Wainwright 2013b; Martin and Feinstein 2014). These species show evidence of pre-mating isolation in the laboratory (West and Kodric-Brown 2015) and are genetically differentiated in sympatry (genome-wide mean *Fst* = 0.14 across 12 million SNPs; (McGirr and Martin 2017)).

We previously identified 31 genomic regions (20 kb) containing SNPs fixed between species (*Fst* = 1), showed signs of a hard selective sweep, and were significantly associated with oral jaw size using multiple genome-wide association mapping approaches (McGirr and Martin 2017). A subset of these fixed SNPs fell within significant QTL explaining 15% of variation in oral jaw size and were near genes annotated for effects on skeletal system development (Martin et al. 2017). Here we use complementary approaches to identify candidate causal variants putatively influencing craniofacial divergence by 1) incorporating transcriptomic data from 122 individuals sampled at three developmental stages (McGirr and Martin 2018; McGirr and Martin 2019), 2) applying genome divergence scans to a much larger sample of whole genomes from San Salvador Island and surrounding Caribbean outgroup populations (increasing n = 37 to 258) aligned to a new high-quality *de novo* genome assembly (Richards et al. 2020), 3) identifying structural variation fixed between species for the first time in this system, and 4) inferring *cis* and *trans* regulatory mechanisms underlying gene expression divergence between species using 12 F1 hybrid transcriptomes. We identified two genes showing *cis*-regulatory divergence that were near just one fixed variant each: a deletion upstream of a gene known to influence skeletal development (*dync2li1*) and a SNP downstream of a novel candidate gene (*pycr3*). Our results highlight the utility of using these closely related species replicated across isolated lake populations as an evolutionary model for craniofacial development and provide highly promising candidate variants for future functional validation experiments.

## Results

### Few fixed variants between young species showing drastic craniofacial divergence

We analyzed whole genome resequencing samples for 258 *Cyprinodon* pupfishes (median coverage = 8×; (Richards et al. 2020)). This included 114 individuals from multiple isolated lake populations on San Salvador Island (33 generalists, 46 molluscivores, and 35 scale-eaters) and 140 outgroup generalist pupfishes from across the Caribbean and North America. Libraries for 150PE Illumina sequencing were generated from DNA extracted from muscle tissue and the resulting reads were mapped to the *C. brontotheroides* reference genome (v 1.0; total sequence length = 1,162,855,435 bp; number of scaffolds = 15,698, scaffold N50, = 32,000,000 bp; L50 = 15 scaffolds; (Richards et al. 2020)). Variants were called using the Genome Analysis Toolkit (GATK v 3.5 (DePristo et al. 2011)) and filtered to include SNPs with a minor allele frequency above 0.05, genotype quality above 20, and sites with greater than 50% genotyping rate across all individuals.

Out of 9.3 million SNPs identified in our dataset, we found a mere 157 SNPs fixed between molluscivore and scale-eater specialist species showing *Fst* = 1 (Fig. 2A; mean genome-wide *Fst* = 0.076). Of these 157 variants, 46 were within 10 kb of 27 genes and none were in coding regions. These 27 genes were enriched for 27 biological processes, including several ontologies describing neuronal development and activity of cell types within bone marrow (Fig. 2B; Table S1).

**Fig. 2.**
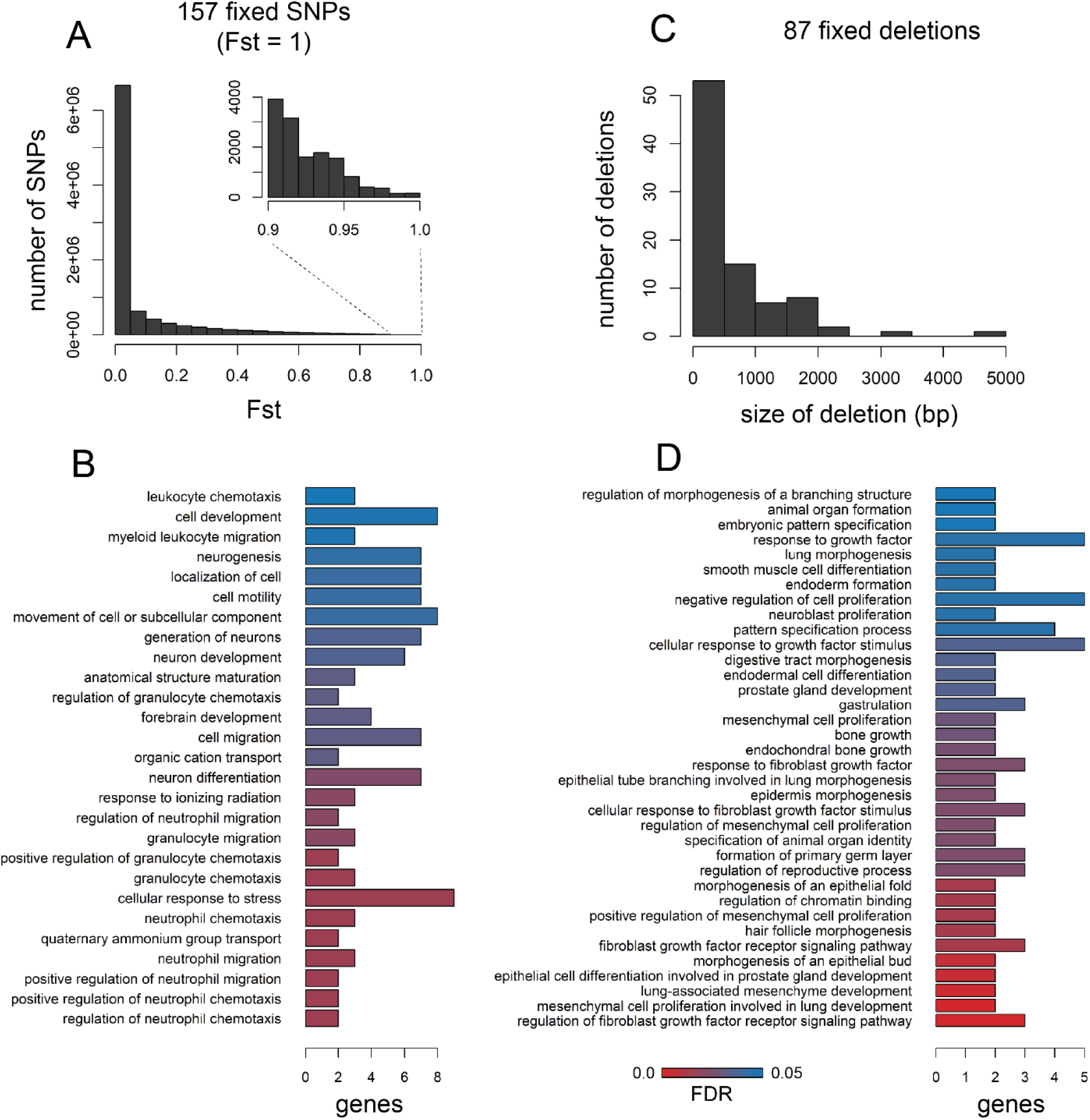
Very few SNPs and structural variants are fixed between trophic specialists. A) Distribution of Weir and Cokerham *Fst* values across 9.3 million SNPs. 157 were fixed between species (*Fst* = 1). B) 46 of the 157 SNPs were located near 27 genes that were enriched for 27 biological processes (FDR < 0.05). C) Size distribution of the 87 deletions are fixed between species out of 80,012 structural variants. D) 34 of the 87 fixed deletions were within 10 kb of 34 genes that were enriched for 36 biological processes.

Structural variants (including insertions, deletions, inversions, translocations, and copy number variants) have been traditionally difficult to detect in non-model systems and ignored by many early whole-genome comparative studies (Stapley et al. 2010; Ho et al. 2019; Wellenreuther et al. 2019). We identified 80,012 structural variants across eight molluscivore and scale-eater individuals using a method that calls variants based on combined evidence from paired-end clustering and split read analysis (Rausch et al. 2012). Just 87 structural variants were fixed between species and, strikingly, all of these variants were deletions fixed in scale-eaters. These deletions ranged in size between 55 bp and 4,703 bp (Fig. 2C). Of these, 34 fixed deletions were near 34 genes (Table S1). Only a single fixed deletion (1,256 bp) was found within a protein coding region, spanning the entire fifth exon of *gpa33* (Fig. 3). The 34 genes near fixed deletions were enriched for 36 biological processes, including ontologies describing bone development, mesenchyme development, fibroblast growth, and digestive tract development (Fig 2D).

**Fig. 3.**
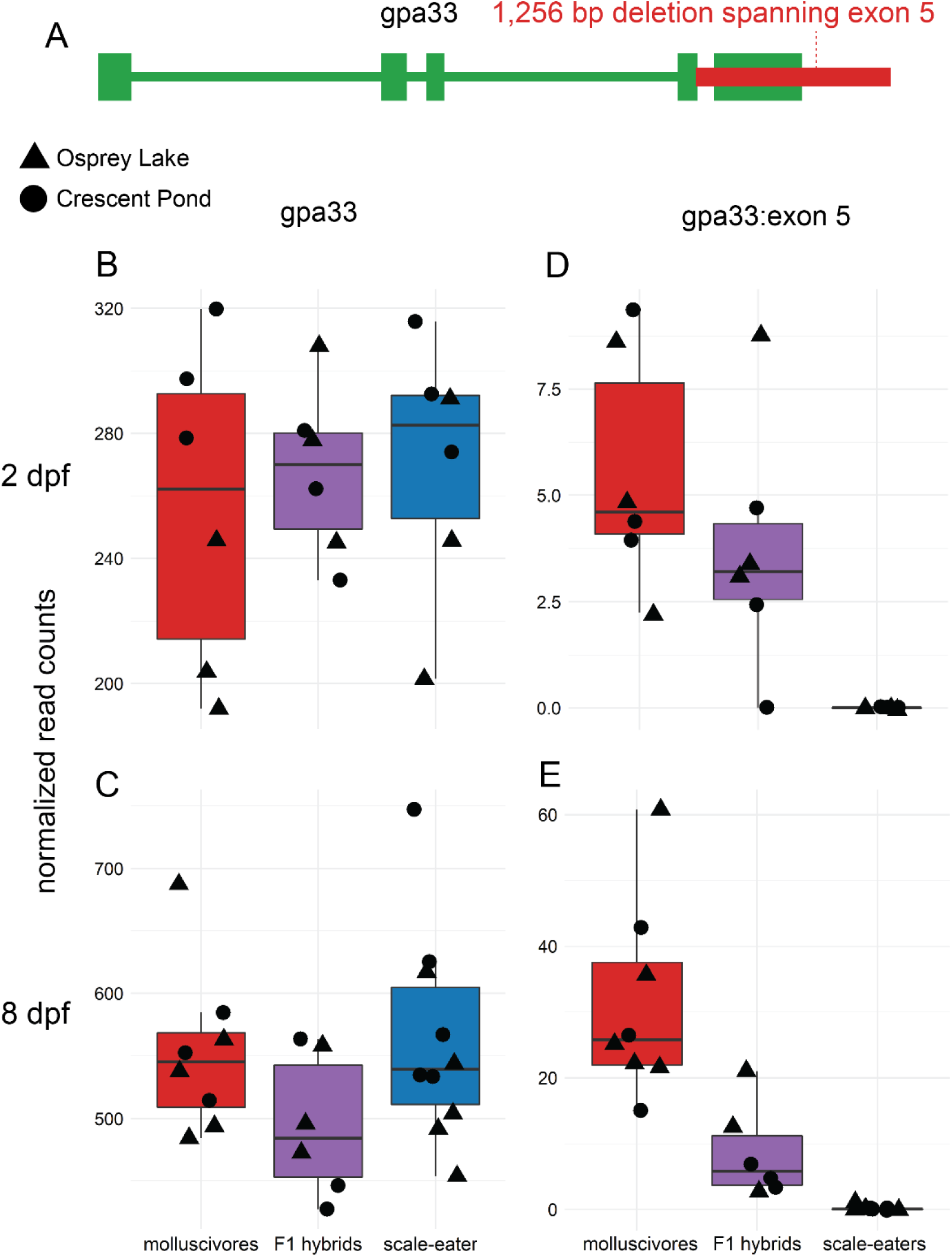
The only fixed variant within a protein coding region is an exon deletion of *gpa33*. A) A 1,256 bp deletion (red) identified by DELLY spans the entire fifth exon of *gpa33* and is fixed in scale-eaters. B and C) The gene is not significantly differentially expressed between molluscivores (red) and scale-eaters (blue) at 2 days post fertilization (dpf) or 8 dpf when considering read counts across all exons (*P* > 0.05). D and E) However, when only considering the fifth exon, scale-eaters show no expression and F1 hybrids (purple) show reduced expression, supporting evidence for the deletion.

Including SNPs and deletions, we found a total of 80 fixed variants within 10 kb of 59 genes (Table S1). Encouragingly, 41 of these genes (70%) also showed high between population nucleotide divergence (*Dxy* > 0.0083 (genome-wide 90^th^ percentile)), strengthening evidence for adaptive divergence at these loci. There are likely many alleles contributing to craniofacial divergence that are segregating between populations of molluscivores and scale-eaters. However, variants with larger effect sizes are predicted to fix faster than variants with smaller effects, especially given short divergence times (Griswold 2006; Yeaman and Whitlock 2011). Thus, these 80 fixed variants provided a promising starting point to identify causal alleles influencing craniofacial phenotype.

### Genes near fixed variants are differentially expressed throughout development

All but one of the 80 variants fixed between species were in non-coding regions, suggesting that they may affect species-specific phenotypes through regulation of nearby genes. In order to identify patterns of gene expression divergence between species, we combined two previous transcriptomic datasets spanning three developmental stages and three San Salvador Island lake populations (McGirr and Martin 2018; McGirr and Martin 2019). F1 offspring were sampled at 2 days post-fertilization (dpf), 8 dpf, and 20 dpf. RNA was extracted from whole body tissue at 2 dpf and 8 dpf; whereas 20 dpf samples were dissected to only extract RNA from craniofacial tissues (Table S2). We used DEseq2 (Love et al. 2014) to contrast gene expression in pairwise comparisons between species grouped by developmental stage (sample sizes for comparisons (molluscivores vs. scale-eaters): 2 dpf = 6 vs. 6, 8 dpf = 8 vs. 10, 20 dpf = 6 vs. 2).

Out of 19,304 genes annotated for the *C. brontotheroides* reference genome, we found 770 (5.93%) significantly differentially expressed at 2 dpf, 1,277 (9.48%) at 8 dpf, and 312 (2.50%) at 20 dpf (Fig. 4A-D). The lower number of genes differentially expressed at 20 dpf likely reflects reduced power to detect expression differences due to the small scale-eater sample size. Nonetheless, we found four genes differentially expressed throughout development at all three stages (*filip1, c1galt1, klhl24*, and *oit3*) and 248 genes were differentially expressed during two of the three stages examined. Of the 59 genes near SNPs or deletions fixed between species, we found 12 differentially expressed during at least one developmental stage (Table 1; Fig. 4E). Two of these genes (*dync2li1* and *pycr3*) were differentially expressed at 2 dpf and 8 dpf.

**Table 1.** Twelve genes differentially expressed between molluscivores and scale-eaters at 2 days post fertilization (dpf), 8 dpf, and/or 20 dpf. Differentially expressed genes showing *cis* regulation showed significant allele-specific expression in F1 hybrids (MBASED *P* < 0.05), while genes showing *trans* regulation did not (MBASED *P* > 0.05). Genes with undetermined regulatory mechanisms underlying expression divergence (NA) had fewer than two informative heterozygous sites within the coding region. MNC = mean normalized counts across all samples. LFC = log2 fold change in expression (positive values indicate higher expression in scale-eaters than molluscivores). *P* = adjusted *P*-value for differential expression.

| gene | mechanism | 2 dpf |  |  | 8 dpf |  |  | 20 dpf |  |  |
| --- | --- | --- | --- | --- | --- | --- | --- | --- | --- | --- |
| | | MNC | LFC | $P$ | MNC | LFC | $P$ | MNC | LFC | $P$ |
| *dync2li1* | *cis* | 96.09 | -0.70 | **3.7E-05** | 34.05 | -1.05 | **5.2E-05** | 23.83 | -1.10 | 1.2E-01 |
| *pycr3* | *cis* | 221.91 | 0.49 | **2.5E-03** | 56.19 | 1.09 | **1.5E-08** | 38.16 | 0.13 | 8.9E-01 |
| *eef1d* | *trans* | 1984.23 | 0.18 | 1.3E-01 | 1076.82 | 0.51 | **8.8E-07** | 1265.39 | 0.08 | 8.9E-01 |
| *washc5* | *trans* | 293.53 | -0.14 | 5.0E-01 | 141.55 | -0.40 | **9.2E-04** | 143.95 | -0.03 | 9.6E-01 |
| *pxk* | *trans* | 205.36 | 0.19 | 2.9E-01 | 183.15 | 0.67 | **1.9E-04** | 120.35 | 0.65 | 7.3E-02 |
| *hint1* | NA | 1719.70 | 0.28 | 2.6E-01 | 824.17 | 0.46 | **9.4E-03** | 336.79 | -1.03 | **9.7E-03** |
| *nsmce2* | NA | 260.89 | -0.48 | **1.4E-04** | 79.51 | -0.44 | 1.5E-02 | 82.97 | -0.80 | 6.1E-02 |
| *gimap2* | NA | 17.46 | 2.14 | **5.5E-04** | 46.44 | 0.04 | 9.5E-01 | 57.94 | 1.89 | 1.6E-02 |
| *cdk5r1* | NA | 106.52 | -0.59 | **3.7E-03** | 292.02 | 0.31 | 9.2E-02 | 7.22 | -1.18 | 3.6E-01 |
| *dph5* | NA | 344.39 | 0.51 | **2.8E-03** | 108.03 | 0.20 | 2.9E-01 | 63.25 | -0.28 | 6.4E-01 |
| *pdhb* | NA | 662.23 | 0.41 | **6.9E-03** | 2359.84 | 0.06 | 8.1E-01 | 680.86 | -0.29 | 5.8E-01 |
| *irf1* | NA | 5.62 | 0.32 | 7.6E-01 | 142.62 | -1.19 | **2.9E-04** | 360.24 | 1.17 | 1.0E-01 |

**Fig. 4.**
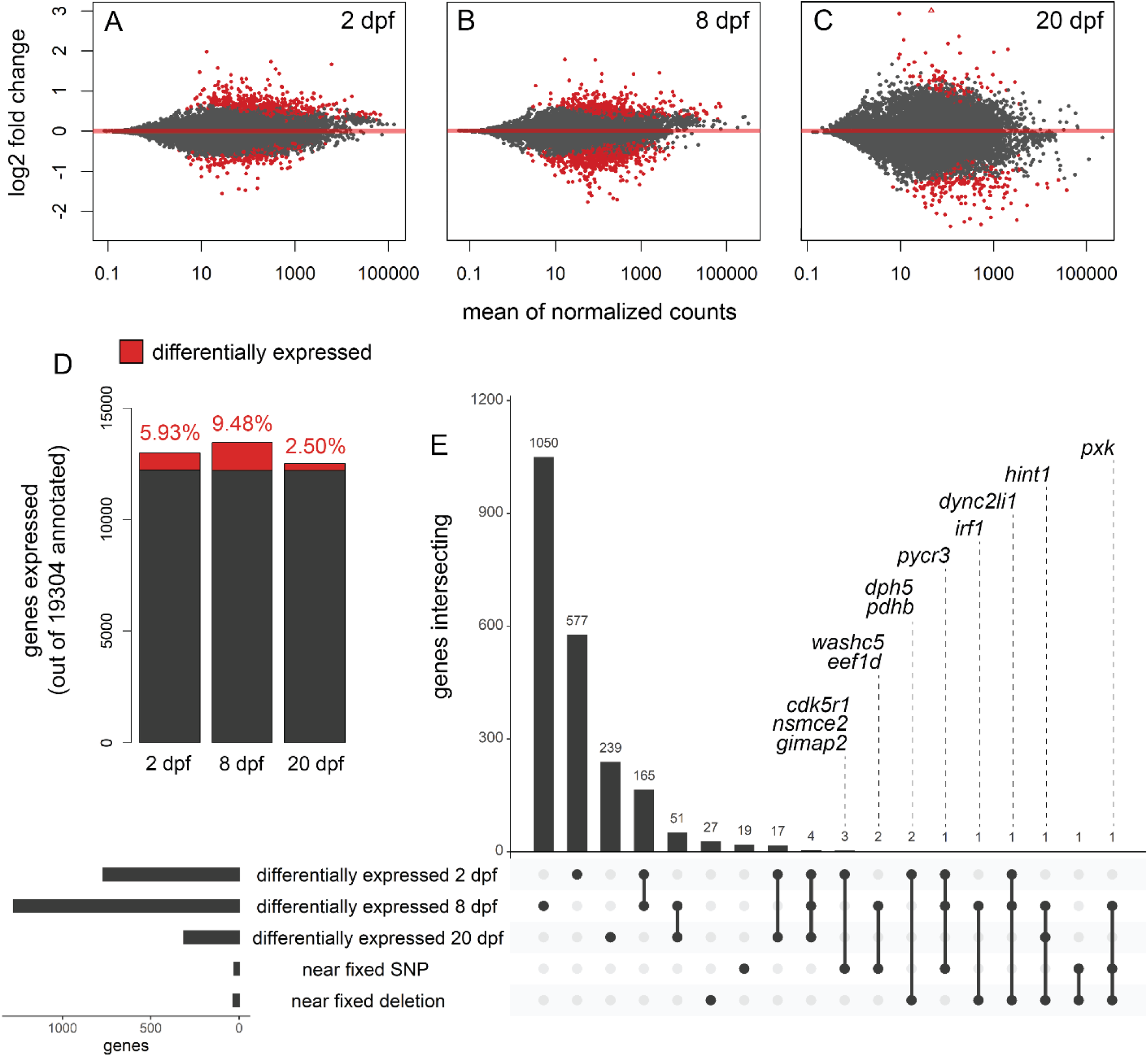
Genes near fixed variants are differentially expressed between species across three developmental stages. Plots show genes differentially expressed (red; *P* < 0.01) between molluscivores and scale-eaters at A) 2 days post fertilization (dpf), B) 8 dpf, and C) 20 dpf. Positive log2 fold changes indicate higher expression in scale-eaters relative to molluscivores. D) Proportion of genes differentially expressed out of the total number of genes expressed across three stages. E) UpSet plot (Conway et al. 2017) showing intersection across five sets: genes differentially expressed at each of the three stages, genes within 10 kb of fixed SNPs, and genes within 10 kb of fixed deletions. The twelve labeled genes were differentially expressed during at least one stage and within 10 kb of fixed variants.

### Fixed variants near genes showing cis-regulatory divergence

In order to determine whether the 12 genes near fixed variants showed differential expression due to *cis*- or *trans*-regulatory divergence, we analyzed expression patterns across 12 F1 hybrid transcriptomes generated from crosses between molluscivores and scale-eaters. The parents used as breeding pairs for these crosses were included in the genomic SNP dataset, allowing us to identify sites that were alternatively homozygous in parents and heterozygous in their F1 hybrids. We measured allele-specific expression (ASE) for genes containing heterozygous sites to identify mechanisms of regulatory divergence (Cowles et al. 2002; Wittkopp et al. 2004). We identified significant ASE using a beta-binomial test comparing the maternal and paternal counts at each gene with the R package MBASED (Mayba et al. 2014). We inferred *cis*-regulatory divergence if a gene was significantly differentially expressed between species and showed significant ASE in all F1 hybrids from a cross at the same developmental stage (Fig. 5A). We inferred *trans*-regulatory divergence if a gene was significantly differentially expressed between species and did not show significant ASE in all F1 hybrids (Fig. 5B).

**Fig. 5.**
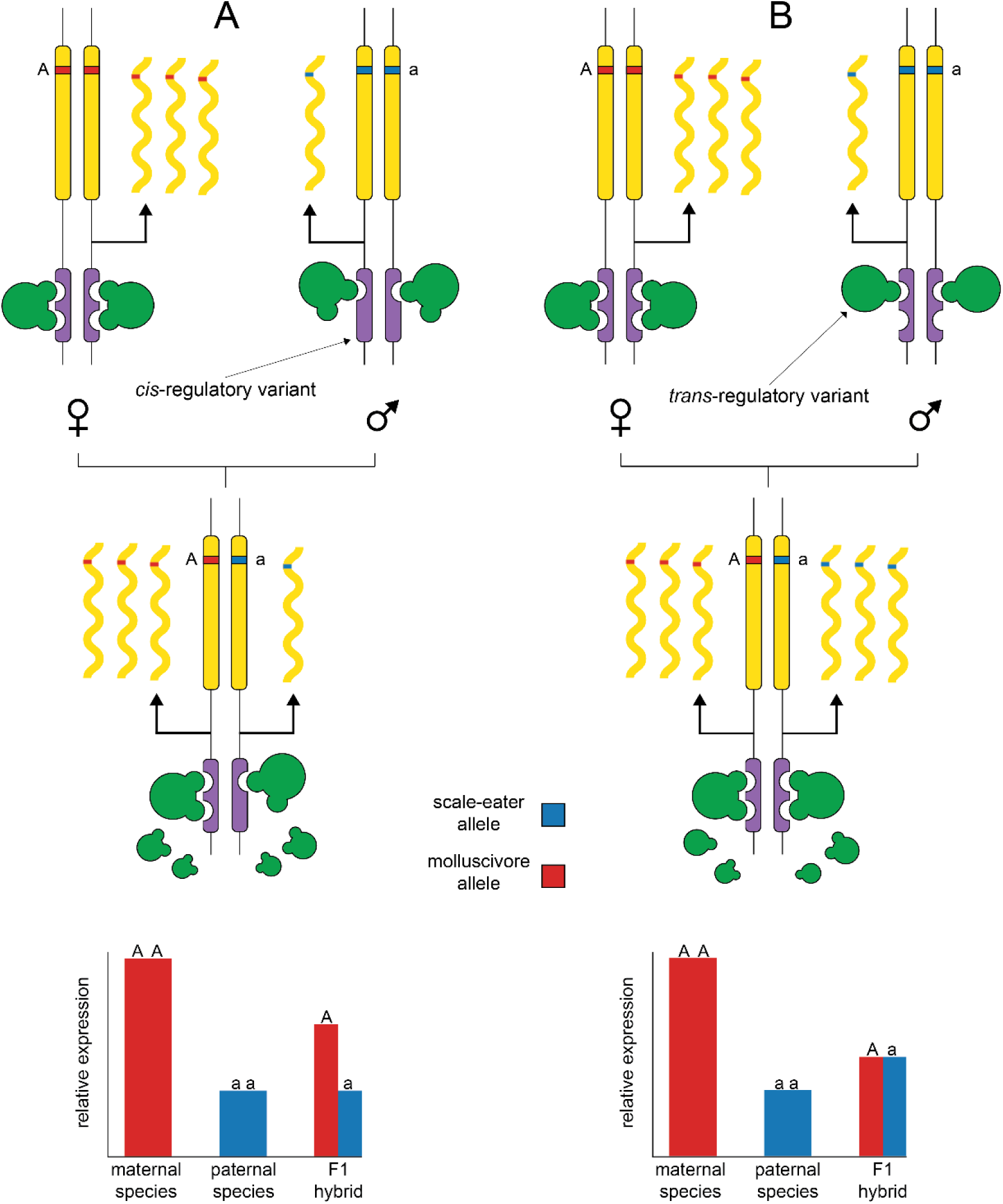
Deciphering between *cis*- and *trans*-regulatory divergence influencing gene expression. Diagrams show protein coding gene regions (yellow) regulated by linked *cis*-acting elements (purple) and *trans*-acting binding proteins (green). In the examples, a female molluscivore is crossed with a male scale-eater to produce an F1 hybrid. The two species are alternatively homozygous for an allele within the coding region of a gene that shows higher expression in the molluscivore than the scale-eater. A) A *cis*-acting variant causing reduced expression results in low expression of the scale-eater allele in the F1 hybrid. B) Lower expression in the scale-eater is caused by a *trans*-acting variant, resulting in similar expression levels of both parental alleles in the F1 hybrid.

Of the 12 genes differentially expressed near fixed variants, five contained at least two informative heterozygous sites that could be used to measure ASE (Fig. 6 and 7). The same two genes that were differentially expressed at 2 dpf and 8 dpf (*dync2li1* and *pycr3*) also showed significant allele specific expression in F1 hybrids at both developmental stages (Fig. 6A and B; MBASED *P* < 0.05). This provided strong evidence that differential regulation of these genes was influenced by nearby fixed variation within putative *cis*-regulatory elements. The three other genes with informative sites (*eef1d, washc5*, and *pxk*) did not show significant ASE, suggesting that *trans-*acting variation may have influenced expression divergence between species for these genes (Fig. 7A-C).

**Fig. 6.**
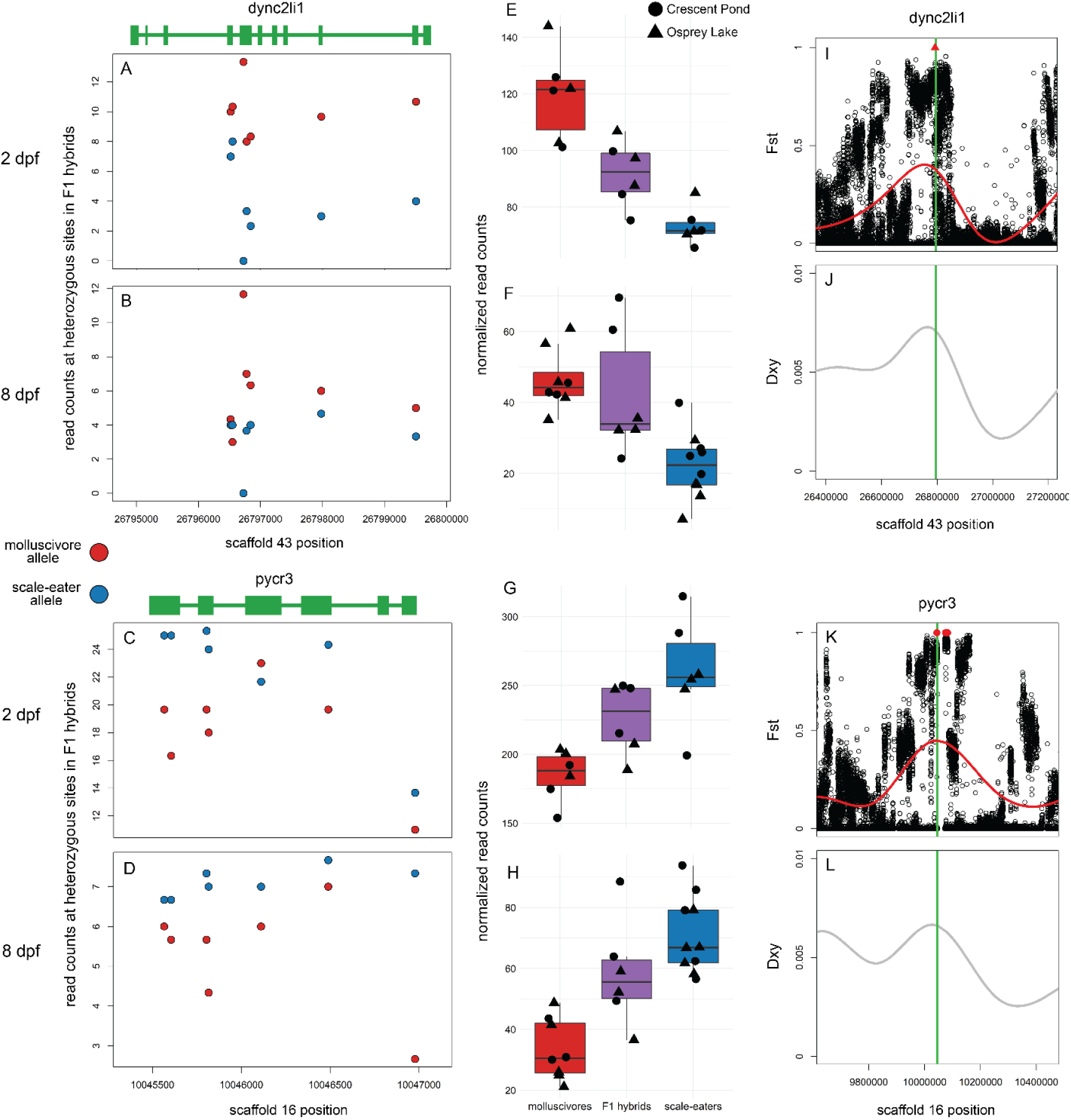
Two genes near fixed variants show *cis*-regulatory divergence between trophic specialists. A-D) Mean counts for reads spanning *dync2li1* and *pycr3* that match parental alleles at heterozygous sites are shown for crosses between Crescent Pond molluscivores (red) and scale-eaters (blue) at 2 dpf (A and C) and 8 dpf (B and D). E-H) Normalized read counts for F1 offspring from Crescent Pond (circles) and Osprey Lake (triangles) crosses. Both genes are differentially expressed between molluscivores (red) and scale-eaters (blue) at both developmental stages (*P* < 0.01) and show additive inheritance in F1 hybrids (purple). For both genes, F1 hybrids show higher expression of alleles derived from the parental species that shows higher gene expression in purebred F1 offspring (MBASED *P* < 0.05), consistent with *cis*- regulatory divergence between species. I-L) Both genes (green lines) are within regions showing high relative genetic differentiation (*Fst*; I and K) and high absolute genetic divergence (*Dxy*; J and L). Red triangle shows fixed deletion. Red points show fixed SNPs (*Fst* = 1).

**Fig. 7.**
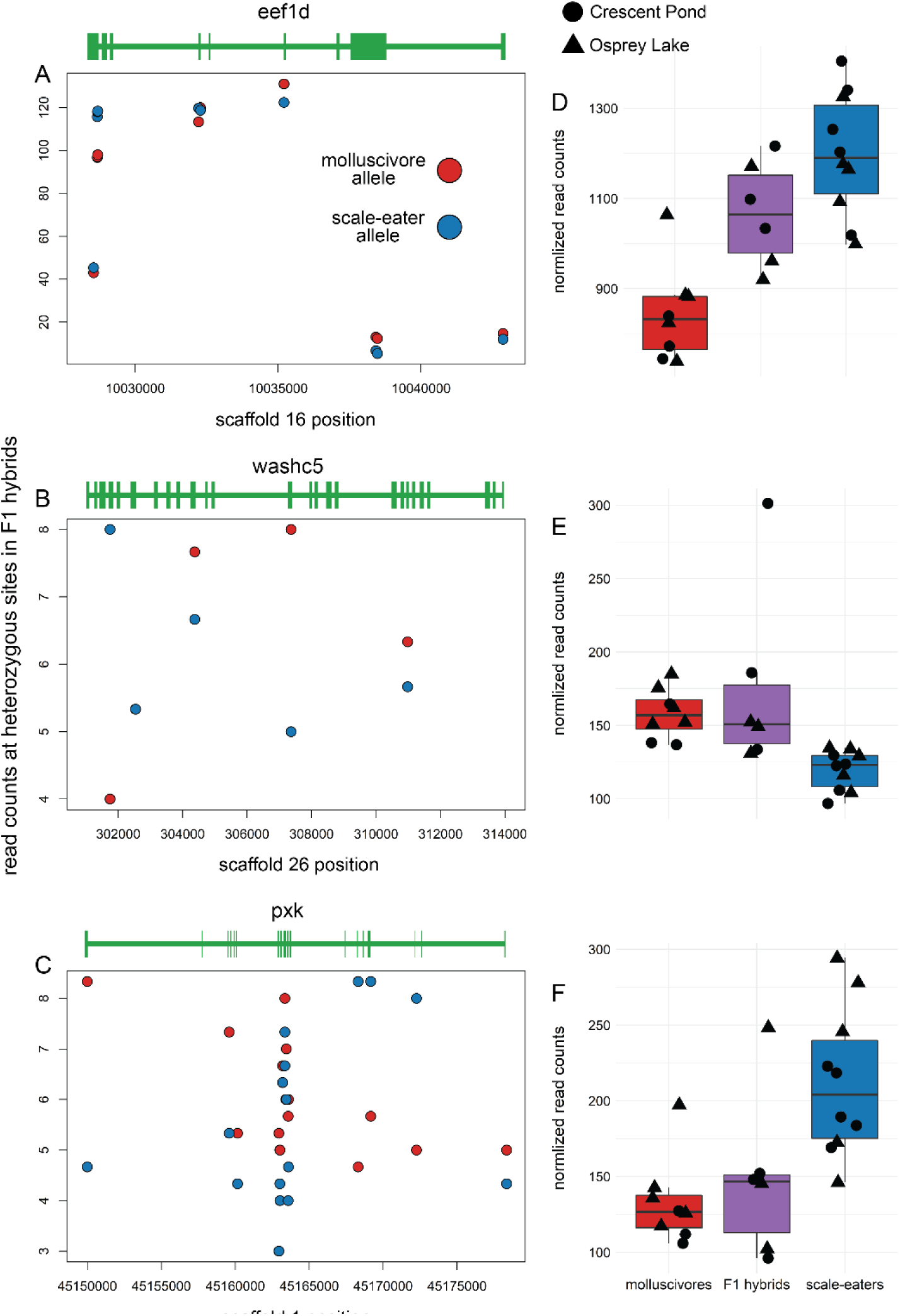
Three genes near fixed variants show *trans*-regulatory divergence between trophic specialists. A-D) Mean counts for reads spanning A) *eef1d*, B) *washc5*, and C) *pxk* that match parental alleles at heterozygous sites are shown for crosses between Crescent Pond molluscivores (red) and scale-eaters (blue) at 8 dpf. D-F) Library size normalized read counts for F1 offspring from Crescent Pond (circles) and Osprey Lake (triangles) crosses. All three genes are differentially expressed between molluscivores (red) and scale-eaters (blue) at 8 dpf. None of these genes showed significant allele-specific expression in F1 hybrids (purple; MBASED, *P* > 0.05), indicating *trans*-regulatory mechanisms underlying expression divergence.

The two genes showing *cis*-regulatory divergence were near just one fixed variant each: a 91 bp deletion located 7,384 bp upstream of *dync2li1* and an A-to-C transversion 1,808 bp downstream of *pycr3* (Fig. 6). The next closest fixed variants were separated by greater than 600 kb and 31 kb, respectively. We searched the JASPAR database (Fornes et al. 2019) to identify potential transcription factor binding sites that could be altered by these candidate *cis*-acting variants. The 91 bp deletion near *dync2li1* contained binding motifs corresponding to seven transcription factors (*nfia, nfix, nfic, znf384, hoxa5, gata1, myb*; Table S3). Two binding motifs spanned the *pycr3* SNP region (*gata2, mzf1*), one of which, *mfz1*, was altered by the alternate allele in scale-eaters. The scale-eater allele created a new potential binding motif matching the transcription factor *plagl2*. Sanger sequencing confirmed the A-to-C transversion near *pycr3* in four additional individuals not included in the whole-genome resequencing dataset (Fig. 8).

**Fig. 8.**
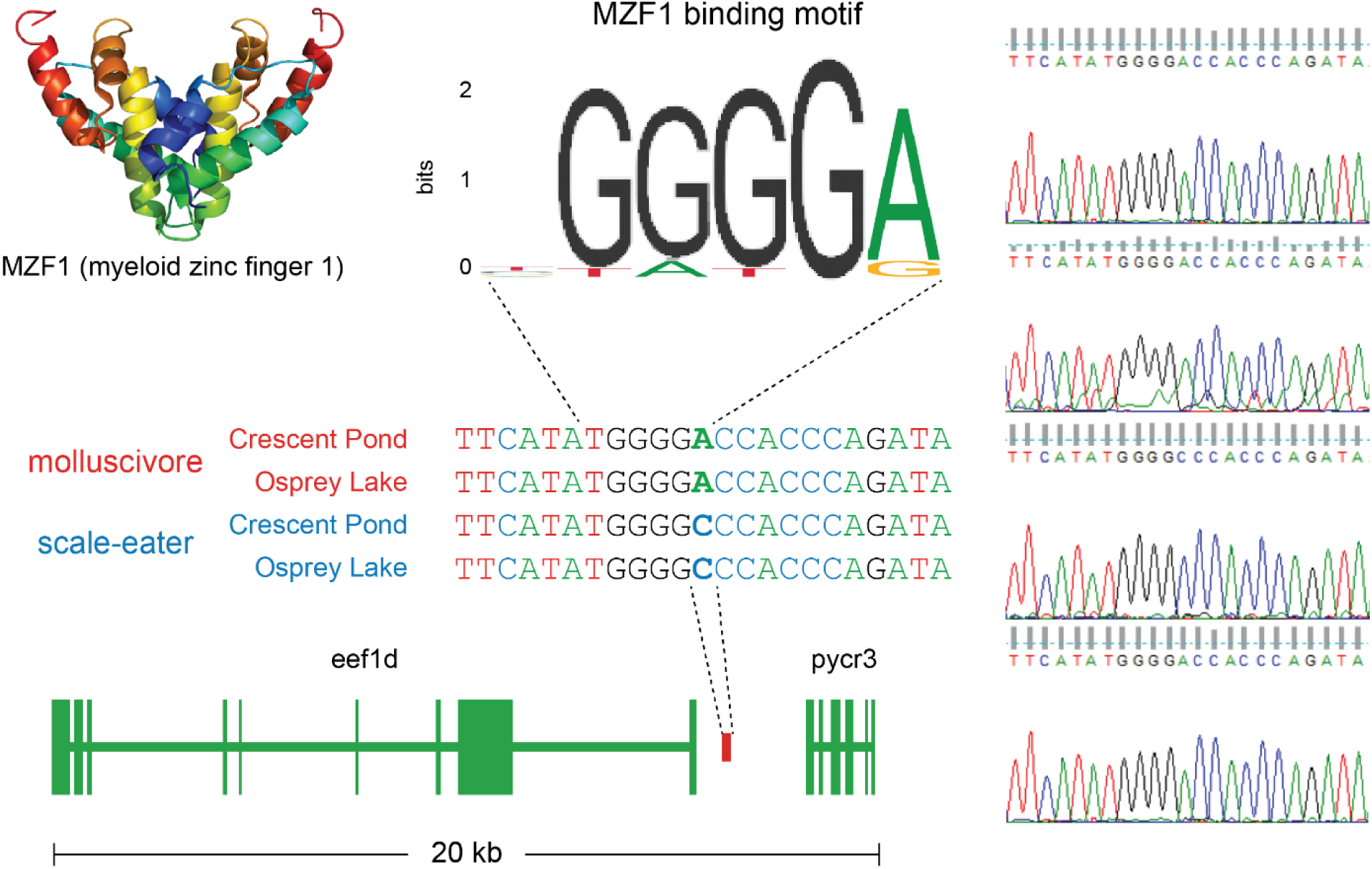
Sanger sequencing confirms fixed SNP that could alter transcription factor binding near *pycr3*. Chromatograms on the right confirm the A-to-C transversion fixed in scale-eaters that falls between *eef1d* (Fig. 7A) and *pycr3* (Fig. 6C-L). The myeloid zinc finger transcription factor binds a motif that matches the molluscivore (JASPAR database matrix ID: MA0056.1), however, the scale-eater allele alters this motif.

## Discussion

Understanding the developmental genetic basis of complex traits by investigating natural variation among closely related species is a powerful complementary approach to traditional genetic screens in model systems. The San Salvador Island *Cyprinodon* pupfish system is a useful evolutionary model for understanding the genetic basis of craniofacial defects and natural diversity given extensive morphological divergence between these young species (Fig. 1). We found just 244 genetic variants fixed between molluscivores and scale-eaters across 9.3 million SNPs and 80,012 structural variants (Fig. 2A and C). Almost all variants were in non-coding regions, with the exception of an exon-spanning deletion (Fig. 3). In support of these variants affecting divergent adaptive phenotypes, 80 variants were near 59 genes that were enriched for developmental functions related to divergent specialist traits (Fig. 2B and D). Furthermore, twelve of these genes were highly differentially expressed between species across three developmental stages (Fig. 4E). By measuring allele-specific expression in F1 hybrids from multiple crosses between species, we found two variants strongly implicated as *cis*-regulatory alleles affecting expression divergence between species: a fixed deletion near *dync2li1* and a fixed SNP near *pycr*3 (Fig. 6).

### Fixed genetic variation underlying trophic specialization

In a previous analysis of SNPs from a smaller whole genome dataset, *dync2li1* was one of 30 candidate genes that showed signs of a hard selective sweep and was significantly associated with variation in jaw size between molluscivores and scale-eaters using multiple genome-wide association mapping approaches (McGirr and Martin 2017). Here we show that a fixed deletion near *dync2li1* may influence expression divergence between species through *cis*-acting regulatory mechanisms. This gene (dynein cytoplasmic 2 light intermediate chain 1) is known to influence skeletal morphology in humans (Kessler et al. 2015; Taylor et al. 2015; Niceta et al. 2018). It is a component of the cytoplasmic dynein 2 complex which is important for intraflagellar transport – the movement of protein particles along the length of eukaryotic cilia (Cole 2003; Pfister et al. 2006). Due to the vital role that cilia play in the transduction of signals in the *hedgehog* pathway and other pathways important for skeletal development, disruptions in dynein complexes cause a variety of skeletal dysplasias collectively termed skeletal ciliopathies (Huber and Cormier-Daire 2012; Taylor et al. 2015). Mutations in *dync2li1* have been linked with ciliopathies that result from abnormal cilia shape and function including Ellis-van Creveld syndrome, Jeune syndrome, and short rib polydactyly syndrome (Kessler et al. 2015; Taylor et al. 2015; Niceta et al. 2018). These disorders are characterized by variable craniofacial malformations including micrognathia (small jaw), hypodontia (tooth absence), and cleft palate (Brueton et al. 1990; Ruiz-Perez and Goodship 2009; Taylor et al. 2015). The discovery of *dync2li1* as a candidate gene influencing differences in oral jaw length between molluscivores and scale-eaters suggests that this system is particularly well-suited as an evolutionary mutant model for clinical phenotypes involving jaw size, such as micrognathia and macrognathia.

We also identified a fixed SNP near the gene *pycr3* (pyrroline-5-carboxylate reductase 3; also denoted *pycrl*) which showed *cis*-regulatory divergence. This gene is not currently known to influence craniofacial phenotypes in humans or other model systems. However, one study investigating gene expression divergence between beef and dairy breed bulls found that *pycr3* was one of the most highly differentially expressed genes in skeletal muscle tissues. The authors found nearly a three-fold difference in expression of *pycr3* between the two bull breeds that are primarily characterized by differences in muscle anatomy (Sadkowski et al. 2009). Similarly, expression changes in this gene may influence skeletal muscle development in specialists species, which differ in the size of their adductor mandibulae muscles (Martin and Wainwright 2011; Hernandez et al. 2018). The A-to-C transversion near *pycr3* could influence differences in expression by altering transcription factor binding. We found that the molluscivore allele matches the binding motif of *mzf1* (myeloid zinc finger 1; Fig. 8), a transcription factor known to influence cell proliferation (Gaboli et al. 2001), whereas the scale-eater allele alters this motif. This type of binding motif analyses has a high sensitivity (*mzf1* is known to bind this motif) but extremely low selectivity (*mzf1* does not bind nearly every occurrence of this motif, which appears 1,430,540 times in the molluscivore reference genome).

While oral jaw size is the primary axis of phenotypic divergence in the San Salvador Island pupfish system, adaptation to divergent niches required changes in a suite of morphological and behavioral phenotypes (St John et al. 2019; St. John et al. 2020). The majority of genes differentially expressed between species were found within whole embryo tissues (Fig. 4A-D), suggesting we should find candidate genes influencing the development of craniofacial phenotypes and other divergent traits. Of the 244 variants fixed between species, the only coding variant was a 1,256 bp deletion that spanned the fifth exon of *gpa33* (glycoprotein A33), which is expressed exclusively in intestinal epithelium (Fig. 3). Knockouts of this gene in mice cause increased hypersensitivity to food allergens and susceptibility to a range of related inflammatory intestinal pathologies (Williams et al. 2015). The gut contents of wild-caught scale-eaters are comprised of 40-51% scales (Martin and Wainwright 2013c). The exon deletion of *gpa33* may play a metabolic role in this unique adaptation that allows scale-eaters to occupy a higher trophic level than molluscivores. Future studies in this system will benefit from sequencing and analyses that target specific tissues and cell types to determine whether candidate variants affect a single phenotype or have pleiotropic effects.

### The effectiveness of Cyprinodon pupfishes for identifying candidate cis-regulatory variants

One major advantage of investigating the genetic basis of craniofacial divergence between molluscivores and scale-eaters is the low amount of genetic divergence between species. Species-specific phenotypes are replicated across multiple isolated lake populations that exhibit substantial ongoing gene flow. This has resulted in small regions of the genome showing strong genetic differentiation, with some regions containing just a single variant fixed between species. Furthermore, a previous study found a significant QTL explaining 15% of variation in oral jaw size and three more potential moderate-effect QTL, suggesting that we may expect to find variants with moderate effects on craniofacial divergence. Thus, we chose to focus our analyses on genes near fixed variation because variants with larger effect sizes are predicted to fix faster than variants with smaller effects, especially given short divergence times (Griswold 2006; Yeaman and Whitlock 2011). The low number of fixed variants dispersed across the genome makes this system relatively unique compared to other systems with similar divergence times (Whiteley et al. 2010; Jones et al. 2012; Martin et al. 2019). Although our approach ignores segregating variation, which likely influences the majority of craniofacial divergence between species, it provides a strategy to identify novel candidate genes like *pycr3* that have not been previously associated with craniofacial development and prioritize such candidates for functional validation experiments.

## Conclusions

Overall, our results highlight the utility of the San Salvador Island pupfish system as an evolutionary mutant model for natural and clinical variation in human craniofacial phenotypes. Similar rapid speciation replicated across many environments can be found in other adaptive radiations (Martin et al. 2019; Martin and Richards 2019), which could also prove useful as evolutionary models for a variety of other human traits. We found that a combination of SNPs and deletions likely contribute to the evolution of highly divergent craniofacial morphology through *cis*-acting effects on the expression of skeletal genes. Future studies will attempt to validate the effect of candidate variation on gene expression and craniofacial development *in vivo*.

## Methods

### Identifying genomic variation fixed between specialists

In order to identify SNPs fixed between molluscivores and scale-eaters, we analyzed whole genome resequencing samples for 258 individuals from across the Caribbean (median coverage = 8×; (Richards et al. 2020)). Briefly, 114 pupfishes from 15 isolated hypersaline lakes and one estuary on San Salvador Island were collected using hand and seine nets between 2011 and 2018. This included 33 generalists, 46 molluscivores, and 35 scale-eaters. Eight of these individuals were bred to generate F1 offspring sampled for RNA sequencing (Table S2). This dataset also included 140 outgroup generalist pupfishes from across the Caribbean and North America, including two individuals belonging to the pupfish radiation in Lake Chichancanab, Mexico, and two individuals from the most closely related outgroups to *Cyprinodon* (*Megupsilon aporus* and *Cualac tessellatus* (Echelle et al. 2005)). Libraries for 150PE Illumina sequencing were generated from DNA extracted from muscle tissue and the resulting reads were mapped to the *C. brontotheroides* reference genome (v 1.0; total sequence length = 1,162,855,435 bp; number of scaffolds = 15,698, scaffold N50, = 32,000,000 bp; L50 = 15 scaffolds; (Richards et al. 2020)). Variants were called using the HaplotypeCaller function of the Genome Analysis Toolkit (GATK v 3.5 (DePristo et al. 2011)) and filtered to include SNPs with a minor allele frequency above 0.05, genotype quality above 20, and sites with greater than 50% genotyping rate across all individuals, resulting in 9.3 million SNPs.

Measuring relative genetic differentiation (*Fst*) between species can point to regions of the genome containing variation affecting divergent phenotypes (Jones et al. 2012; Poelstra et al. 2014; Lamichhaney et al. 2015). However, *Fst* is dependent on the many potential forces acting to reduce within-population nucleotide diversity, including selective sweeps, purifying selection, background selection, and low recombination rates (Noor and Bennett 2009; Cruickshank and Hahn 2014). Measuring between-population divergence (*Dxy*) can help distinguish between these possibilities because nucleotide divergence between species increases at loci under different selective regimes (Nachman and Payseur 2012; Cruickshank and Hahn 2014; Irwin et al. 2016). We measured *Fst* between species with vcftools (v. 0.1.15; weir-fst-pop function) and identified fixed SNPs (*Fst* = 1). We also measured *Fst* and *Dxy* in 10 kb windows using the python script popGenWindows.py created by Simon Martin (github.com/simonhmartin/genomics_general; (Martin et al. 2013)).

We identified structural variation (insertions, deletions, inversions, translocations, and copy number variants) fixed between specialist species with DELLY (v 0.8.1; (Rausch et al. 2012)). Unlike GATK HaplotypeCaller which is limited to identifying structural variants smaller than half the length of read size (DePristo et al. 2011), DELLY can identify small variants in addition to variants larger than 300 bp using paired-end clustering and split read analysis. We used DELLY to identify structural variants across eight whole genomes from the breeding pairs used to generate F1 hybrid RNA samples (Four scale-eaters from two lake populations and four molluscivores from the same two lake populations; Table S2). First, we trimmed reads using Trim Galore (v. 4.4, Babraham Bioinformatics), aligned them to the *C. brontotheroides* reference genome with the Burrows-Wheeler Alignment Tool (v 0.7.12; (Li and Durbin 2011), and removed duplicate reads from the resulting .bam files with Picard MarkDuplicates (broadinstitute.github.io/picard). Second, we called variants with DELLY by comparing an individual of one species with all individuals of the other species, resulting in eight variant call files. Third, we identified structural variants fixed between species that were shared across all eight files, in which all molluscivores showed the reference allele and all scale-eaters showed the same alternate allele.

### Transcriptomic sequencing, alignment, and variant discovery

Our transcriptomic dataset included 50 libraries from 122 individuals sampled across three early developmental stages (Table S2; (McGirr and Martin 2018; McGirr and Martin 2019)). Breeding pairs used to generate F1 hybrids and purebred F1 offspring were collected from three hypersaline lakes on San Salvador Island: Crescent Pond, Osprey Lake, and Little Lake. For purebred crosses, we collected F1 embryos from breeding tanks containing multiple breeding pairs from a single lake population. For F1 hybrid samples, we crossed a single individual of one species with a single individual of another species from the same lake population.

RNA was extracted from samples collected two days after fertilization (2 dpf) eight days after fertilization (8 dpf), and 17-20 days after fertilization (20 dpf) using RNeasy Mini Kits (Qiagen catalog #74104). For samples collected at 2 dpf, we pooled 5 embryos together and pulverized them in a 1.5 ml Eppendorf tube using a plastic pestle washed with RNase Away (Molecular BioProducts). We used the same extraction method for samples collected at 8 dpf but did not pool larvae and prepared a library for each individual separately. We dissected samples collected at 20 dpf to isolate tissues from the anterior craniofacial region containing the dentary, angular articular, maxilla, premaxilla, palatine, and associated craniofacial connective tissues using fine-tipped tweezers washed with RNase AWAY. The earlier developmental stages are described as stage 23 (2 dpf) and 34 (8 dpf) in a recent embryonic staging series of *C. variegatus* (Lencer and McCune 2018). The 2 dpf stage is comparable to the Early Pharyngula Period of zebrafish, when multipotent neural crest cells have begun migrating to pharyngeal arches that will form the oral jaws and most other craniofacial structures (Schilling and Kimmel 1994; Furutani-Seiki and Wittbrodt 2004; Lencer et al. 2017). Embryos usually hatch six to ten days post fertilization, with similar variation in hatch times among species (Lencer et al. 2017; McGirr and Martin 2018). While some cranial elements are ossified prior to hatching, the skull is largely cartilaginous at 8 dpf and ossified by 20 dpf (Lencer and McCune 2018). All samples were reared in breeding tanks at 25–27°C, 10–15 ppt salinity, pH 8.3, and fed a mix of commercial pellet foods and frozen foods.

Methods for total mRNA sequencing were previously described (McGirr and Martin 2018; McGirr and Martin 2019). Briefly, 2 dpf and 8 dpf libraries were prepared using TruSeq stranded mRNA kits and sequenced on 3 lanes of Illumina 150 PE Hiseq4000 at the Vincent J. Coates Genomic Sequencing Center (McGirr and Martin 2019). All 20 dpf libraries were prepared using the KAPA stranded mRNA-seq kit (KAPA Biosystems 2016) at the High Throughput Genomic Sequencing Facility at UNC Chapel Hill and sequenced on one lane of Illumina 150PE Hiseq4000 (McGirr and Martin 2018). We filtered raw reads using Trim Galore (v. 4.4, Babraham Bioinformatics) to remove Illumina adaptors and low-quality reads (mean Phred score < 20) and mapped 122,090,823 filtered reads to the *C. brontotheroides* reference genome (Richards et al. 2020) using the RNAseq aligner STAR with default parameters (v. 2.5 (Dobin et al. 2013)). We assessed mapping and read quality using MultiQC (Ewels et al. 2016) and quantified the number of duplicate reads and the median percent GC content of mapped reads for each sample using RSeQC (Wang et al. 2012). Although all reads were mapped to a molluscivore reference genome, we did not find a significant difference between species in the proportion of reads uniquely mapped with STAR (Fig. S1 A; Student’s t-test, *P* = 0.061). Additionally, we did not find a difference between species in the proportion of multimapped reads, GC content of reads, or number of duplicate reads (Fig. S1 B-D; Student’s t-test, *P* > 0.05).

We used GATK HaplotypeCaller function to call SNPs across 50 quality filtered transcriptomes. We refined SNPs using the allele-specific software WASP (v. 0.3.3) to correct for potential mapping biases that would influence tests of allele-specific expression (ASE; (Van De Geijn et al. 2015)). WASP identified reads that overlapped SNPs in the initial .bam files and re-mapped those reads after swapping the genotype for the alternate allele. Reads that failed to map to exactly the same location were discarded. We re-mapped unbiased reads to create our final .bam files used for differential expression analyses. Finally, we re-called SNPs using unbiased .bam files for allele specific expression analyses.

### Differential expression analyses

We used the featureCounts function of the Rsubread package (Liao et al. 2014) requiring paired-end and reverse stranded options to generate read counts across 19,304 genes and 156,743 exons annotated for the molluscivore (*C. brontotheroides*) reference genome (Richards et al. 2020). We used DESeq2 (v. 3.5 (Love et al. 2014)) to normalize raw read counts for library size and perform principal component analyses, and identify differentially expressed genes. DESeq2 fits negative binomial generalized linear models for each gene across samples to test the null hypothesis that the fold change in gene expression between two groups is zero. Significant differential expression between groups was determined with Wald tests by comparing normalized posterior log fold change estimates and correcting for multiple testing using the Benjamini–Hochberg procedure with a false discovery rate of 0.01 (Benjamini and Hochberg 1995).

We constructed a DESeqDataSet object in R using a multi-factor design that accounted for variance in F1 read counts influenced by parental population origin and sequencing date (design = ∼sequencing_date + parental_breeding_pair_populations). Next, we used a variance stabilizing transformation on normalized counts and performed a principal component analysis to visualize the major axes of variation in 2 dpf, 8 dpf, and 20 dpf samples (Fig. S2). We contrasted gene expression in pairwise comparisons between species grouped by developmental stage (sample sizes for comparisons (molluscivores vs. scale-eaters): 2 dpf = 6 vs. 6, 8 dpf = 8 vs. 10, 20 dpf = 6 vs. 2). We used plyranges (v. 1.6.5; (Lee et al. 2019)) to determine if genetic variation fixed between species fell within 10 kb of significantly differentially expressed genes (> 10 kb from the start of the first exon and <10 kb from the end of the last exon).

### Allele specific expression analyses

It is possible to identify mechanisms of gene expression divergence between parental species by bringing *cis* elements from both parents together in the same *trans* environment in F1 hybrids and quantifying allele specific expression (ASE) of parental alleles at heterozygous sites (Fig. 5; (Cowles et al. 2002; Wittkopp et al. 2004)). A gene that is differentially expressed between parental species that also shows allele specific expression biased toward one parental allele is expected to result from *cis*-regulatory divergence. A gene that is differentially expressed between parental species that does not show ASE in F1 hybrids is expected to result from *trans*-regulatory divergence. After identifying genes differentially expressed between species that were also near fixed variants, we wanted to test whether those genes showed signs of *cis*-regulatory divergence. This would indicate that fixed variation likely contributed to expression divergence between species.

Our SNP dataset included every parent used to generate F1 hybrids between populations (*n* = 8). We used the GATK VariantsToTable function (DePristo et al. 2011) to output genotypes across 9.3 million SNPs for each parent and overlapped these sites with the variant sites identified in F1 hybrid transcriptomes. We used python scripts (github.com/joemcgirr/fishfASE) to identify SNPs that were alternatively homozygous in breeding pairs and heterozygous in their F1 offspring. We counted reads across heterozygous sites using ASEReadCounter (-minDepth 20 --minMappingQuality 10 --minBaseQuality 20 -drf DuplicateRead) and matched read counts to maternal and paternal alleles. For genes that were differentially expressed between species that were near fixed variants, we identified significant ASE using a beta-binomial test comparing the maternal and paternal counts at each gene with the R package MBASED (Mayba et al. 2014). For each F1 hybrid sample, we performed a 1-sample analysis with MBASED using default parameters run for 1,000,000 simulations to determine whether genes showed significant ASE in hybrids (*P* < 0.05).

For genes within 10 kb of variants fixed between species, we inferred *cis*-regulatory divergence if a gene was significantly differentially expressed between species (DeSeq2 *P* < 0.01) and showed significant ASE in all F1 hybrids from a cross at the same developmental stage (MBASED; *P* < 0.05). We inferred *trans*-regulatory divergence if a gene was significantly differentially expressed between species (DeSeq2 *P* < 0.01) and did not show significant ASE in all F1 hybrids (MBASED; *P* > 0.05; Fig. 5). We required that genes had at least two informative SNPs with ≥10× coverage to assign *cis*- or *trans-* regulatory divergence.

We most likely underestimated the number of fixed variants acting as *cis*-regulatory alleles influencing expression divergence between species. First, variants could affect expression at a developmental stage not included in our sampling. Second, because our approach to identify regulatory mechanisms underlying expression divergence relies on F1 hybrid expression, the advantage of having low genetic variation between species is counterbalanced by the disadvantage of limited heterozygosity within coding regions that provide informative sites to estimate allele-specific expression. Out of twelve genes near fixed variants that were differentially expressed between species, only five contained more than one heterozygous informative site to assign *cis*- or *trans*- regulatory divergence. Third, some of the genes that we classified as *trans*-regulated showed low overall levels of expression, reducing our power to detect significant differences in expression levels of parental alleles. Fourth, we required that genes show allele-specific expression across the entire coding region to assign *cis*-regulatory divergence, which ignored the possibility of alleles affecting the expression of specific transcript isoforms. For these reasons, our estimation of *cis*-regulatory divergence was highly conservative but still provided promising candidate genes for future study.

### Gene ontology enrichment and transcription factor binding site analyses

We performed gene ontology (GO) enrichment analyses for genes near candidate adaptive variants using ShinyGo v.0.51 (Ge et al. 2019). The *C. brontotheroides* reference genome was annotated using MAKER, a genome annotation pipeline that annotates genes, transcripts, and proteins (Cantarel et al. 2008). Gene symbols for orthologs identified by this pipeline largely match human gene symbols. Thus, we searched for enrichment across biological process ontologies curated for human gene functions.

We searched the JASPAR database (Fornes et al. 2019) to identify whether fixed variation near genes showing *cis*-regulatory divergence altered potential transcription factor binding sites. We generated fasta sequences for the molluscivore containing the variant site and 20 bp on either end of the site and searched across all 1011 predicted vertebrate binding motifs in the database using a 95% relative profile score threshold. We then preformed the same analysis for scale-eater fasta sequences containing the alternate allele.

### Genotyping fixed variants

In order to confirm the genotypes of putative *cis*-acting variants, we performed Sanger sequencing on four additional individuals that were not included in our whole-genome dataset. We extracted DNA from muscle tissue using DNeasy Blood and Tissue kits (Qiagen, Inc.) from two molluscivores and two scale-eaters (one sample from Crescent Pond and one from Osprey Lake for both species). We designed primers targeting the regions containing variation fixed between species near the two genes showing evidence for *cis*-regulatory divergence (*pycr3* and *dync2li1*) using the NCBI primer design tool (Ye et al. 2012). We designed primers targeting a 446 bp region containing the SNP fixed between species (scaffold: HiC_scaffold_16 ; position: 1,0043,644) that was 1,808 bp downstream of *pycr3* (forward: 5′- ACCATTCCAGAAGACAAAAAGCG-3′; reverse: 5′-GGCCCTATATATGGGATGCACAA- 3′). Sequences were amplified with PCR using New England BioLabs *Taq* polymerase (no. 0141705) and dNTP solution (no. 0861609) and Sanger sequencing was performed at Eton Bioscience Inc. (Research Triangle Park, North Carolina). Aligning the resulting sequences using the Clustal Omega Multiple Sequence Alignment Tool (Madeira et al. 2019)) confirmed the A-to-C transversion in scale-eaters (Fig. 8). We designed two additional primer sets targeting the deletion region near *dync2li1* (scaffold: HiC_scaffold_43 ; position: 26,792,380-26,792,471). While both primer sets amplified the sequence in molluscivore samples (not shown), we were unable to amplify this region in scale-eaters, potentially due to high polymorphism in this region.

## Acknowledgements

This study was funded by the University of North Carolina at Chapel Hill, the Miller Institute for Basic Research in the Sciences, NSF CAREER Award 1749764, and NIH/NIDCR R01 DE027052 to CHM. Research was also supported by a Graduate Research Fellowship from the Triangle Center for Evolutionary Medicine and an SSE Rosemary Grant Travel Award to JAM. We thank Daniel Matute, Emilie Richards, Michelle St. John, Bryan Reatini, and Sara Suzuki for valuable discussion; The Vincent J. Coates Genomics Sequencing Laboratory at the University of California, Berkeley for performing RNA library prep and Illumina sequencing; the Gerace Research Centre for logistics; and the Bahamian government BEST Commission for permission to conduct this research.

## Data Accessibility

All transcriptomic raw sequence reads are available as zipped fastq files on the NCBI BioProject database. Accession: PRJNA391309. Title: Craniofacial divergence in Caribbean Pupfishes. All R and Python scripts used for pipelines are available on Github (github.com/joemcgirr/fishfASE).

## Author Contributions

JAM wrote the manuscript, extracted the RNA samples, and conducted all bioinformatic and population genetic analyses. Both authors contributed to the conception and development of the ideas and revision of the manuscript.

