## Supplemental Tables S1-3 and Figures S1-2 for "Conspicuous candidate alleles point to *cis*-regulatory divergence underlying rapidly evolving craniofacial phenotypes"

**Table S1.** Protein coding genes near 157 SNPs and 87 deletions fixed between molluscivores and scale-eaters (within 10 kb of the first or last exon).

| near fixed SNP | near fixed deletion |
|---|---|
| *cckar* | *acat2* |
| *cdc14ab* | *acvr1c* |
| *cdk5r1* | *adra2db* |
| *cxcr1* | *cckar* |
| *dapk2* | *cep170* |
| *derl1* | *col12a1* |
| *dysf* | *ctnnb1* |
| *eef1d* | *dph5* |
| *fev* | *dync2li1* |
| *gimap2* | *eef1a1* |
| *nabp1* | *fam219a* |
| *nat14* | *fgfr2* |
| *nsmce2* | *gm11992* |
| *polg* | *gpa33* |
| *prpf4b* | *hint1* |
| *pxk* | *hlf* |
| *pycr3* | *hlh-13* |
| *sbk2* | *irf1* |
| *sgk1* | *kcnq5* |
| *slc25a29* | *lyrm7* |
| *slc38a2* | *med25* |
| *vrtn* | *mprip* |
| *washc5* | *ncl1* |
| *wdr78* | *odf3l2* |
| *wnt7b* | *pdhb* |
| *zhx2* | *pld5* |
| *znf628* | *pxk* |
| | *rabgap1l* |
| | *sh3pxd2a* |
| | *shisa2* |
| | *slc30a7* |
| | *u2af2* |
| | *upp2* |
| | *znf865* |

**Table S2.** Cross design used to produce RNA sequencing libraries for F1 offspring sampled at 2 days post fertilization (dpf), 8 dpf, and 20 dpf. CP = Crescent Pond, OL = Osprey Lake, and LL = Little Lake.

| Mother | Father | Stage | Libraries | F1 |
|---|---|---|---|---|
| CP molluscivore | CP scale-eater | 2 dpf | 3 | hybrid |
| OL scale-eater | OL molluscivore | 2 dpf | 3 | hybrid |
| CP molluscivore | CP scale-eater | 8 dpf | 3 | hybrid |
| OL scale-eater | OL molluscivore | 8 dpf | 3 | hybrid |
| CP molluscivore | CP molluscivore | 2 dpf | 3 | purebred |
| CP scale-eater | CP scale-eater | 2 dpf | 3 | purebred |
| OL molluscivore | OL molluscivore | 2 dpf | 3 | purebred |
| OL scale-eater | OL scale-eater | 2 dpf | 3 | purebred |
| CP molluscivore | CP molluscivore | 8 dpf | 3 | purebred |
| CP scale-eater | CP scale-eater | 8 dpf | 5 | purebred |
| OL molluscivore | OL molluscivore | 8 dpf | 5 | purebred |
| OL scale-eater | OL scale-eater | 8 dpf | 5 | purebred |
| CP molluscivore | CP molluscivore | 20 dpf | 3 | purebred |
| CP scale-eater | CP scale-eater | 20 dpf | 2 | purebred |
| LL molluscivore | LL molluscivore | 20 dpf | 3 | purebred |

**Table S3.** Predicted transcription factor binding sites (JASPAR database) altered by genetic variants fixed between species.

| gene region | allele | transcription factor | matrix ID | binding sequence | relative profile score |
|---|---|---|---|---|---|
| dync2li1 | reference | NFIC | MA0161.1 | TTGGCA | 1.00000 |
| dync2li1 | reference | NFIA | MA0670.1 | AATGCCAAGT | 0.98703 |
| dync2li1 | reference | NFIX | MA0671.1 | AATGCCAAG | 0.98551 |
| dync2li1 | reference | ZNF384 | MA1125.1 | TCAGAAAAAAAA | 0.96526 |
| dync2li1 | reference | HOXA5 | MA0158.1 | CTGTAATT | 0.96148 |
| dync2li1 | reference | Gata1 | MA0035.1 | TGATGC | 0.95591 |
| dync2li1 | reference | MYB | MA0100.3 | CACAACTGGC | 0.95232 |
| dync2li1 | reference | Prrx2 | MA0075.1 | AATTA | 1.00000 |
| dync2li1 | reference | Stat5a | MA1624.1 | GTTCCAAGAATT | 0.98454 |
| dync2li1 | alternate | Prrx2 | MA0075.1 | AATTA | 1.00000 |
| dync2li1 | alternate | Stat5a | MA1624.1 | GTTCCAAGAATT | 0.98454 |
| pycr | reference | GATA2 | MA0036.1 | AGATA | 0.97565 |
| pycr | reference | MZF1 | MA0056.1 | GGGGGA | 0.96199 |
| pycr | alternate | PLAGL2 | MA1548.1 | TGGGCCCCCA | 0.98454 |
| pycr | alternate | GATA2 | MA0036.1 | AGATA | 0.97565 |

**Fig. S1.** Quality control measures for 50 RNAseq libraries. We did not find a difference between scale-eaters and mollusciores in A) the proportion of reads uniquely mapped to the molluscivore reference genome (Student's t-test, *P* = 0.061), B) the proportion of multimapped reads (Student's t-test, *P* = 0.14) ,C) the median GC content of aligned reads (Student's t-test, *P* = 0.22), or D) the number of duplicate reads (Student's t-test, *P* = 0.05).

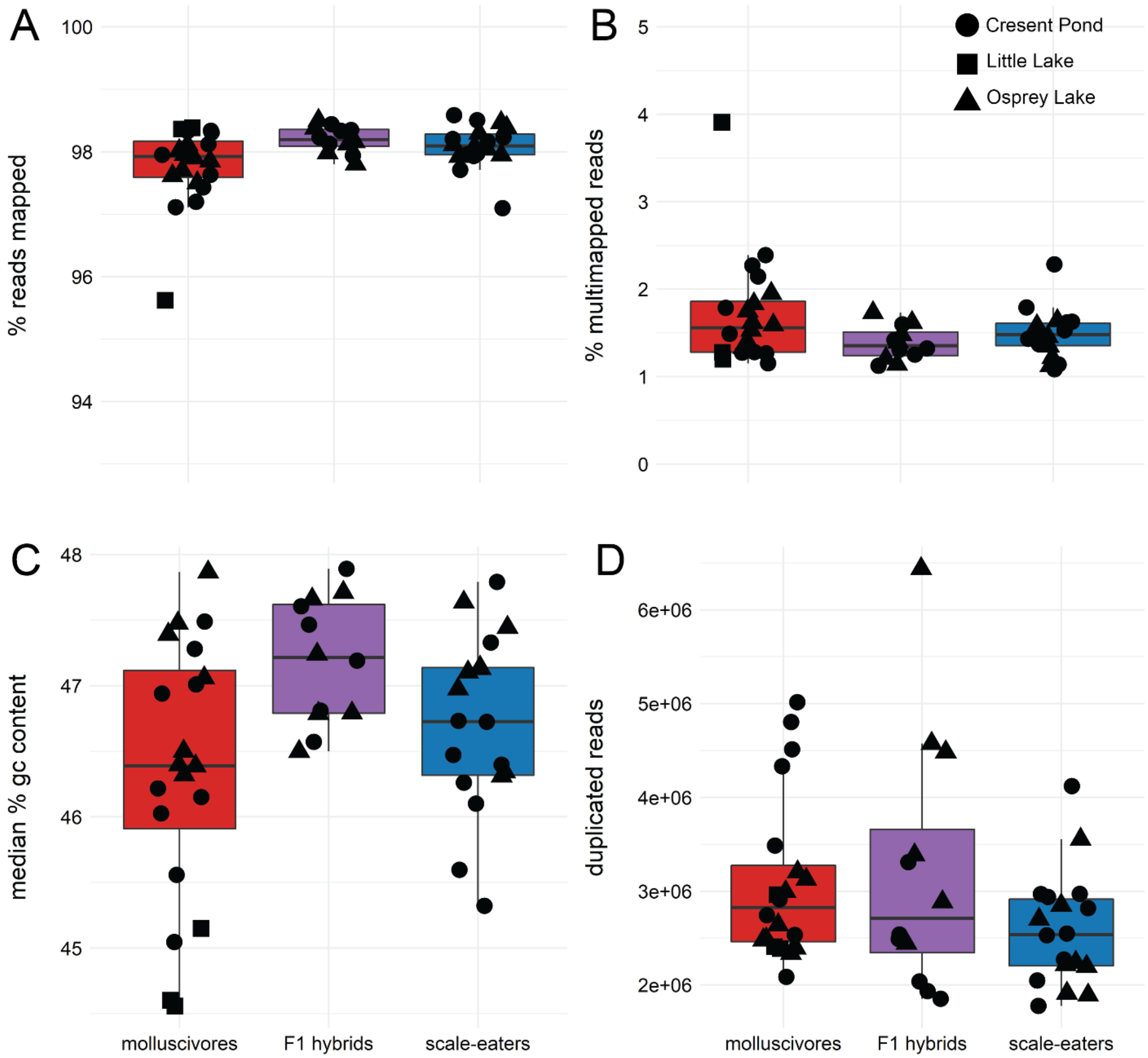

**Fig. S2.** Principal component analysis for 50 transcriptomes showing first two axes accounting for a combined 91% of the total variation in read counts normalized for library size.

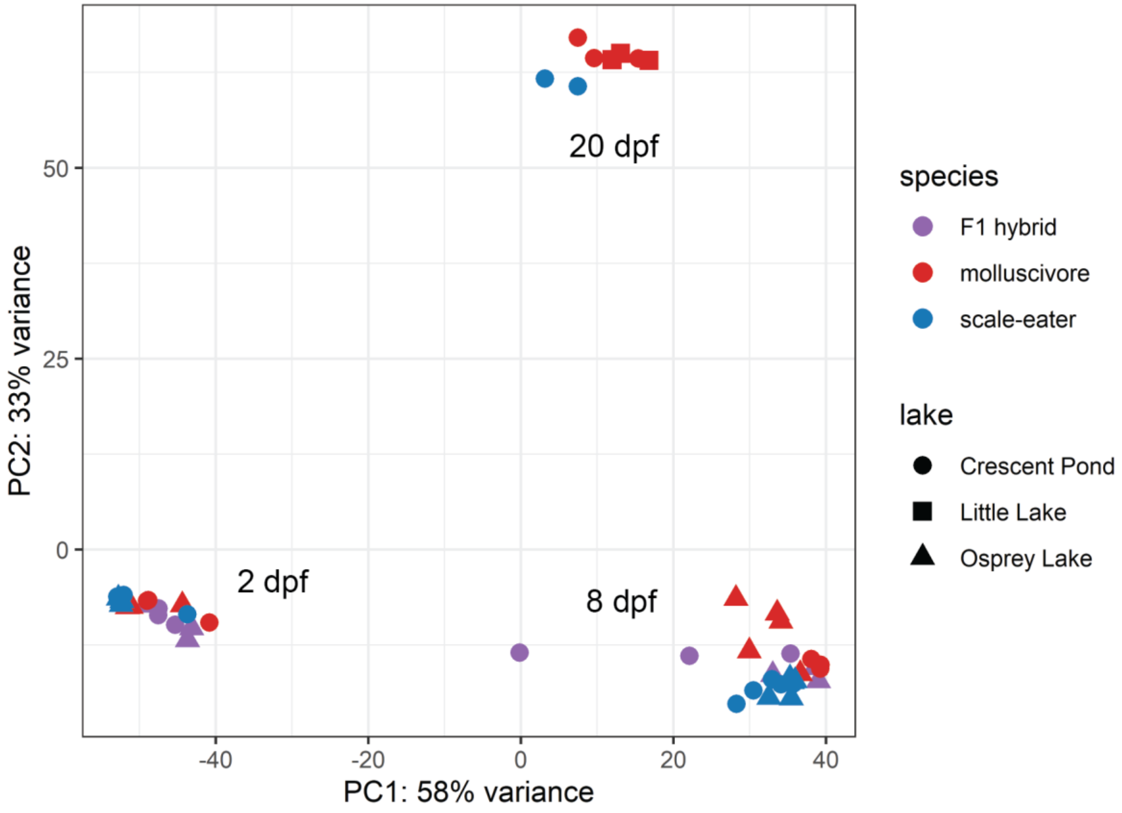
